# Shared but temporally distinct neural representations support semantic matching across word and picture formats: evidence from EEG decoding and temporal generalization analyses

**DOI:** 10.64898/2026.07.06.736728

**Authors:** Ying Xin, Huashuai Xu, Fengyu Cong, Weiqi He, Guanghui Zhang

**Author notes:** **Correspondence:** Guanghui Zhang and Weiqi He, Institute of Psychological and Brain Sciences, Liaoning Normal University, No. 850, Huanghe Road, Shahekou District, Dalian 116029, China. (Guanghui Zhang); (Weiqi He).

## Abstract

Audiovisual semantic matching can be achieved using either written words or pictures, yet whether these formats engage shared semantic matching representations with similar temporal dynamics remains unclear. We recorded electroencephalography from 27 participants while they performed audiovisual semantic matching tasks in which spoken words were paired with either written words or pictures. Stimuli included both natural and man-made objects. Time-resolved multivariate pattern analyses (MVPA or decoding), cross-decoding, and temporal generalization analyses were used to characterize the temporal dynamics of semantic processing. Reliable decoding of matching versus mismatching judgments emerged in both word and picture conditions. Decoding onset that significant above chance level occurred earlier for written words than for pictures and cross-decoding analyses revealed successful generalization between word and picture formats. Temporal generalization analyses further demonstrated distinct representational dynamics across formats, with word processing characterized by predominantly time-specific neural representations and picture processing showing more sustained and temporally stable representations. In addition, matching-related discrimination emerged earlier for natural objects than for man-made objects across both formats. The results suggest that speech-word matching shows earlier neural evidence of audiovisual alignment than speech-picture matching, potentially reflecting differences in how auditory linguistic input is integrated with visual information across representational formats.

## 1. Introduction

In everyday life, people frequently match spoken language with information presented in other visual formats. For example, after hearing the word “apple,” one can quickly determine whether a subsequently presented written word or picture refers to the same object. This ability relies on the integration of auditory input with visual information, ultimately supporting semantic matching and object recognition (Chen & Spence, 2013; Ghuman et al., 2026; Potter & Faulconer, 1975; Theios & Amrhein, 1989). Although both written words and pictures convey conceptual information (Dirani & Pylkkänen, 2024; Ganis et al., 1996; Kutas & Federmeier, 2011), they differ in representational format and may therefore engage distinct neural processing pathways.

One potential source of this difference lies in how auditory and visual information are aligned across representational formats. Written words share linguistic codes with spoken words and may therefore be more readily aligned with auditory word representations. In contrast, pictures require the extraction and organization of visual object information before this visual information can be integrated with the auditory word (Dirani & Pylkkänen, 2024; Gao et al., 2026; Kutas & Federmeier, 2011). Consequently, speech–word matching may show earlier neural evidence of audiovisual alignment than speech–picture matching (Dirani & Pylkkänen, 2023; Ghazaryan et al., 2023; Rahimi et al., 2022). Importantly, such differences should not be interpreted as evidence that written words necessarily provide faster semantic access than pictures, but rather as reflecting differences in the integration of auditory and visual information across formats.

Recent advances in multivariate pattern analysis (MVPA; decoding) provide a powerful framework for addressing this question. Compared with traditional event-related potential (ERP) analyses, MVPA is more sensitive to distributed patterns of neural activity and allows the characterization of how neural representations emerge, evolve, and generalize over time (Carrasco et al., 2024; Grootswagers et al., 2017; Li Calzi et al., 2025; Zhang et al., 2025). For example, some previous studies comparing words and pictures have shown that semantic category information can be decoded from both stimulus formats and that they evoke partially overlapping neural representations despite differences in perceptual processing (Bezsudnova et al., 2024; Dirani & Pylkkänen, 2024). MVPA has also been used to investigate relationships between visual and auditory processing. For example, visual and auditory object categorization has been shown to involve both common and modality-specific representational dynamics (Iamshchinina et al., 2022). Additionally, cross-modal decoding studies have further demonstrated the integration of auditory and visual information during associative memory retrieval and letter-speech sound processing, suggesting that information from different modalities can converge onto common representational codes (Maack et al., 2025; Xu et al., 2025). However, most prior work has examined either word–picture relationships within visual modalities or unimodal semantic categorization. Whether speech-word and speech-picture semantic matching recruit shared neural representations, and whether these representations emerge with similar or distinct temporal profiles, remains unknown.

In the present study, we conducted time-resolved decoding and temporal generalization analyses to investigate the temporal dynamics of semantic matching across different representational formats. Specifically, we recorded EEG signals while participants performed two cross-modal semantic matching tasks. That is, a speech-word matching task and a speech-picture matching task. In each task, participants heard a spoken word and judged whether it matched a simultaneously presented written word or picture (e.g., the spoken word “apple” paired with the written word or picture “apple” or “orange”). In addition to decoding congruency semantic, we also examined whether neural activity patterns retained information about representational format (word vs. picture) even when semantic matching status was held constant. Furthermore, previous theories of semantic cognition suggest that natural and man-made objects differ in their underlying feature structure. Natural objects tend to be characterized by highly shared perceptual features, whereas man-made objects rely relatively more on functional, contextual, and associative information for conceptual representation (Hoffman et al., 2018; Ralph et al., 2017; Tyler et al., 2000). Therefore, we also examined whether object category (natural objects such as “apple” vs. artificial objects such as “candle”) modulates the temporal emergence of semantic matching representations. This design allowed us to directly compare the time course of semantic processing across modality (word vs. picture) and category (natural vs. artificial) within a unified framework.

We hypothesized that (1) speech-word and speech-picture matching would both elicit reliable decoding of matching versus mismatching conditions, reflecting the emergence of shared semantic matching representations across modalities; (2) decoding onset for speech–word matching would occur earlier than speech–picture matching, indicating earlier emergence of semantic matching representations in the speech–word condition. Such differences may reflect more efficient alignment between spoken and written linguistic representations rather than differences in semantic access alone; (3) cross-decoding analyses would reveal shared neural matching representations between tasks, and we expected picture processing to exhibit broader temporal generalization than word processing; and (4) natural objects would show earlier decoding onset than artificial objects across both word and picture conditions, reflecting differences in perceptual and conceptual processing demands between object categories.

## 2. Methods

### 2.1 Participants

We used G*Power 3.1 to perform an a priori power analysis, which indicated that a minimum sample size of 23 participants would be sufficient to detect the expected effects, assuming a large-to-medium effect size (Cohen’s d = 0.80) and a significance level of α = 0.05.

Twenty-eight healthy students from Liaoning Normal University (12 males; 18–25 years old, mean age = 21.6 ± 1.98 years) participated in the experiment. One participant was excluded because their mean behavioral accuracy was below 60%, leaving 27 participants for all EEG analyses. All participants were right-handed, had normal or corrected-to-normal vision, and reported no history of neurological or psychiatric disorders. Written informed consent was obtained prior to participation. The study was approved by the Institutional Review Board of Liaoning Normal University (No.: LL2026153) and conducted in accordance with the Declaration of Helsinki.

### 2.2 Experimental material

The experiment was programmed and conducted in MATLAB R2022a. Visual stimuli were presented using Psychtoolbox (Version 3.0.19). The experimental materials consisted of pictures, written words, and auditory stimuli. Picture stimuli were selected from the databases of Krautz et al. (2022) and Snodgrass and Vanderwart (1980). Following screening for picture complexity, Chinese name frequency, and name complexity, 52 images were retained for the experiment. Half of the stimuli represented natural objects and half represented artificial objects. All pictures were normalized for luminance using the SHINE_color toolbox (Dal Ben, 2023; Willenbockel et al., 2010).

Corresponding word stimuli consisted of two-character Chinese names referring to the depicted objects. Word stimuli were presented in black SimHei font (250 pt) on a white background, with font type and size kept constant across all conditions.

Auditory stimuli were delivered through CANYON-G6 over-ear headphones. Speech recordings were generated using the Ondoku text-to-speech platform:https://ondoku3.com/zh-hans/, employing two male and two female synthetic voices. All auditory stimuli were approximately 550 ms in duration, matched for loudness, and processed using identical parameters. That is, the audio files were saved in .wav format with a sampling rate of 24 kHz, 32-bit depth, and mono channel configuration.

Visual stimuli were displayed on a 23.8-inch Lenovo LED monitor with a resolution of 1920 × 1080 pixels and a refresh rate of 60 Hz. Participants were seated approximately 57 cm from the screen. At this viewing distance, each visual stimulus subtended a visual angle of approximately 13.8°.

### 2.3 Procedure

On each trial, one of four stimulus types (matching word, mismatching word, matching picture, or mismatching picture) was randomly presented at the center of the screen. Participants were instructed to place their hands on a standard QWERTY keyboard, with the left index finger resting on the “F” key and the right index finger on the “J” key. For half of the participants, the “F” key was assigned to matching stimuli and the “J” key to mismatching stimuli. For the remaining participants, the response mapping was reversed. Word in Chinese and picture stimuli were presented with equal probability (1:1), as were matching and mismatching trials. In total, 208 trials were included in the experiment.

Before the formal experiment, participants completed a practice session consisting of 20 trials to familiarize themselves with the task. During the formal experiment, each of the four conditions was presented 52 times, yielding 208 trials that were randomly distributed across four blocks.

Each trial began with a fixation cross presented for a randomly jittered duration of 200–500 ms, followed by the presentation of the visual stimulus (Figure 1). The auditory stimulus was presented 150 ms before the onset of the visual stimulus, following previous studies showing that this stimulus onset asynchrony helps align the timing of auditory and visual semantic activation (Chen & Spence, 2011), thereby facilitating audiovisual semantic matching. The auditory stimulus lasted approximately 550 ms. The visual stimulus remained on the screen for a maximum of 1500 ms or until a response was made. Participants were instructed to determine whether the auditory and visual stimuli were semantically congruent and to respond within 1500 ms. This was followed by a blank screen with a jittered duration of 800–1000 ms.

**Figure 1.**
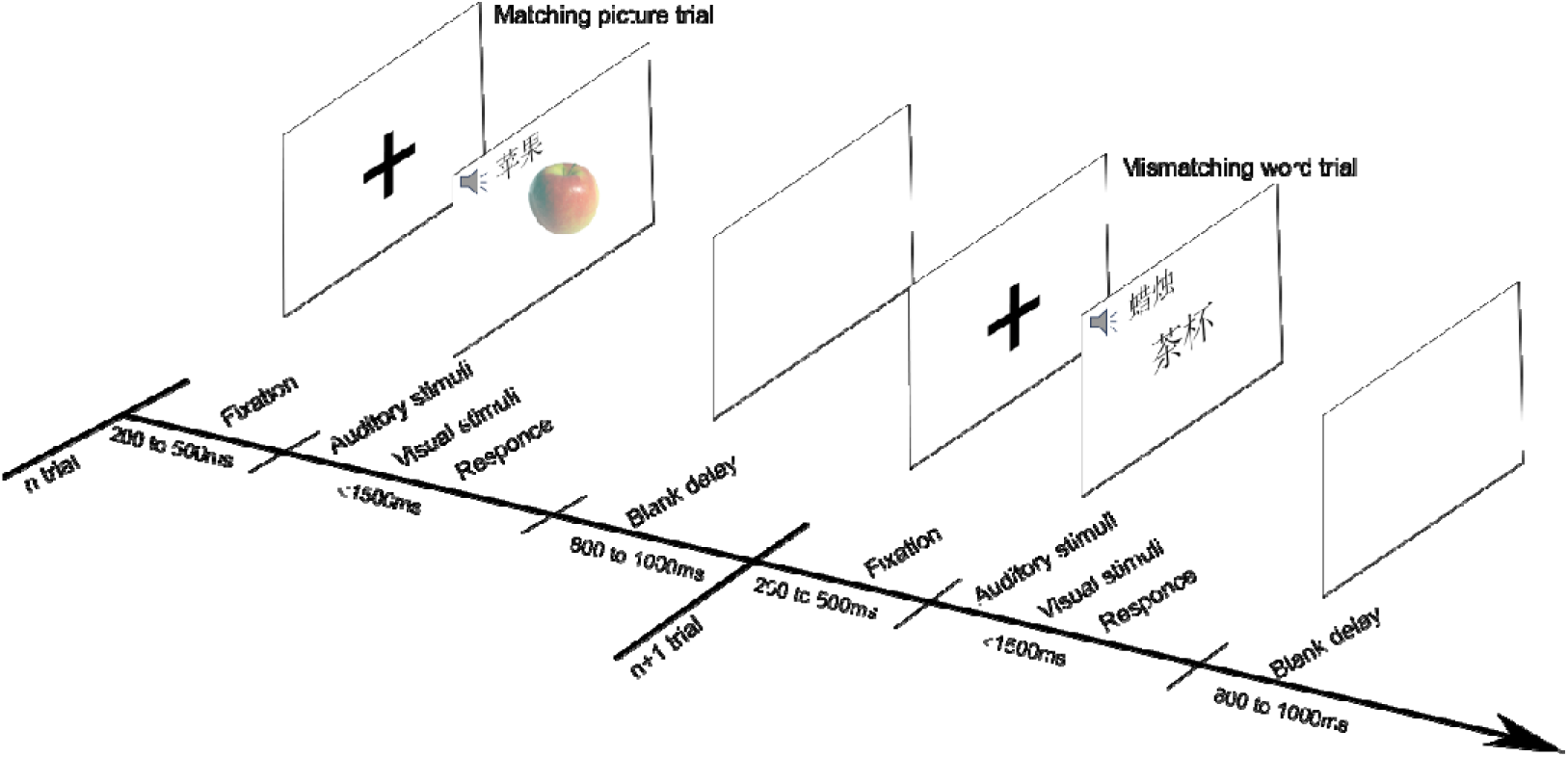
Examples of the stimuli used in this experiment. Each trial began with a fixation cross presented for 200-500 milliseconds, followed by a visual stimulus. The auditory stimulus onset preceded the visual stimulus by 150 milliseconds, and its duration was approximately 550 milliseconds. The visual stimulus was presented for a maximum of 1500 milliseconds. This was followed by a blank screen with a jittered duration ranging from 800 to 1000 milliseconds. Participants were asked whether the auditory stimulus matched with the meaning of visual stimulus within 1500 ms.

### 2.4 EEG recording and preprocessing

EEG data were acquired using a NeuroScan SynAmps amplifier (SynAmps2TM Mode 8050, NeuroScan) at a sampling rate of 1000 Hz. Recordings were obtained from 64 scalp electrodes positioned according to the extended International 10–20 system, including FPz/1/2, AF3/4, Fz/1/2/3/4/5/6/7/8/11/12, FT11/12, FCz/1/2/3/4/5/6/, T7/8, TP7/8, C1/2/3/4/5/6, CPz/1/2/3/4/5/6, M1/2, Pz/1/2/3/4/5/6/7/8, POz/3/4/7/8, Oz/1/2. Electrooculogram (EOG) activity was monitored using electrodes placed lateral to the eyes and beneath the left eye. Electrode impedances were maintained below 5 kΩ throughout data acquisition.

EEG preprocessing was conducted offline in MATLAB 2023a environment using EEGLAB V2023a (Delorme & Makeig, 2004) and ERPLAB V13.0 (Lopez-Calderon & Luck, 2014) toolboxes. Continuous EEG signals were re-referenced to the average of the left and right mastoid electrodes (M1 and M2). To attenuate line noise, a 48–52 Hz notch filter was applied. Data were subsequently band-pass filtered between 0.1 and 30 Hz using a noncausal Butterworth filter with a roll-off of 48 dB/octave (Zhang, Garrett, & Luck, 2024a, 2024b). Ocular artifacts related to blinks and eye movements were corrected using independent component analysis implemented with an optimized procedure as used in prior studies (Dimigen, 2020; S. J. Luck, 2022; Zhang, Garrett, Simmons, et al., 2024). Noisy electrodes were interpolated using spherical spline algorithm implemented in ERPLAB (Buss & Fillmore, 2001).

The continuous data were segmented into epochs extending from -200 to 800 ms relative to stimulus onset. Baseline correction was performed using the prestimulus interval from -200 to 0 ms. The previous studies demonstrated that artifact rejection does not uniformly improve decoding performance and may sometimes reduce classification accuracy (Bae & Luck, 2018; Kessler et al., 2025; Zhang & Luck, 2025), we therefore didnot exclude any trials from further analysis. To ensure that the neural signals reflected successful task performance, only trials associated with correct behavioral responses were included in the subsequent analyses.

### 2.5 SVM-based decoding analysis

In this study, we conducted four independent decoding analyses. First, we decoded whether the presented stimuli were matching or mismatching trials separately for words and pictures. Second, we performed cross-decoding analyses between word and picture categories. Specifically, we first trained the decoder on matching and mismatching trials from the word category and then tested it on matching and mismatching trials from the picture format, and vice versa. Third, to examine semantic differences between words and pictures, we separately decoded word and picture data for matching trials and mismatching trials. Fourth, to examine whether natural objects and man-made objects shared similar semantic matching representations, we decoded matching versus mismatching trials separately within the natural and man-made objects for words and pictures.

We conducted decoding analyses independently at each time point for each participant, and each of the four decoding analyses. For all binary classification analyses, support vector machine (SVM) classifiers were implemented in MATLAB using the fitcsvm() function. Classification performance was quantified with the predict() function by calculating decoding accuracy on the held-out test data. We employed the default regularization parameter (BoxConstraint =1) to balance decoding accuracy with generalization as previous studies (Zhang, Wang, Winsler, et al., 2026; Zhang, Wang, Xin, et al., 2026). Note that the EEG data were resampled to 50 Hz prior to decoding analysis, and EEG signals from 60 channels were included in the decoding procedure.

To reduce overfitting and accommodate the relatively small number of trials, we employed a within-subject N-fold cross-validation procedure with 3 folds, following approaches used in previous ERP decoding studies (Hong et al., 2020; Song et al., 2025; Zhang et al., 2024; Zhang & Luck, 2025). For each stimulus condition, trials were randomly partitioned into 3 subsets without replacement, and trials within each subset were averaged to create ERP average (Hou et al., 2025, 2026; Sarrett & Toscano, 2024; Xin et al., 2026). During each cross-validation iteration, two averaged ERP sets were used for classifier training and the remaining set was reserved for testing. This procedure was repeated until every ERP set had served as the test set once. To improve the stability and reliability of decoding estimates, the complete analysis pipeline was repeated 100 times with different random trial assignments.

To assess the temporal stability of the neural representations identified by the time-resolved decoding analyses, we performed temporal generalization analyses following the procedure described by King and Dehaene (2014). Specifically, for each independent decoding analysis, classifiers trained at each time point were tested across all remaining time points, yielding a temporal generalization matrix (TGM). This approach allows the examination of whether information encoded at one moment is reinstated or maintained at later stages of processing. Generalization extending beyond the diagonal is typically interpreted as evidence for stable neural representations that persist over time, whereas decoding confined mainly to the diagonal reflects transient neural states characterized by continuously changing representational formats (King & Dehaene, 2014).

### 2.6 Statistical analysis for decoding performance

For statistical evaluation of decoding performance and TGM, we tested whether decoding accuracy exceeded the theoretical chance level of 50%, as commonly performed in previous ERP decoding studies (Bae & Chen, 2024; Bae & Luck, 2018). Statistical significance was assessed using a nonparametric cluster-based permutation approach similar to cluster-level mass univariate analyses widely adopted in EEG research (Maris & Oostenveld, 2007). Specifically, one-sample t-tests were conducted at each post-stimulus time point to determine whether decoding accuracy was significantly above chance for the first three decoding analyses. Because below-chance decoding is generally not interpretable for SVM classifiers, one-tailed tests were used. Adjacent time points reaching p < 0.05 were combined into temporal clusters, and the summed t-values within each cluster were used as the cluster-level statistic (t-mass). The observed cluster statistics were then compared with a null distribution generated through permutation testing to control for multiple comparisons across time. Statistical significance was determined using a family-wise alpha threshold of 0.05.

### 2.7 Statistical analysis for behavioral data

We performed a 2-way repeated measures analysis of variance (rmANOVA) on behavioral data [i.e., reaction time (RT), accuracy], with visual format (word vs. picture) and matching (matching vs. mismatching) as within-subject factors. We used Greenhouse-Geisser corrections and post hoc comparisons with the Bonferroni correction if necessary. Partial eta squared (*η_p_*^2^) was used as indices of effect size.

## 3. Results

### 3.1 Behavioral results

For behavioral accuracy (see Figure 2a), the results revealed a significant main effect for visual format [F_(1,26)_ =9.16, *p* = 0.006, *η_p_*^2^ = 0.26], with higher accuracy observed in the word visual format compared to the picture visual format. In contrast, neither the main effect of matching factor [F_(1,26)_ =0.12, *p* = 0.73, *η_p_*^2^ = 0.005] nor the interaction between visual format and matching factors [F_(1,26)_ <0.01, *p* = 1.0, *η_p_*^2^ < 0.001] reached statistical significance.

**Figure 2.**
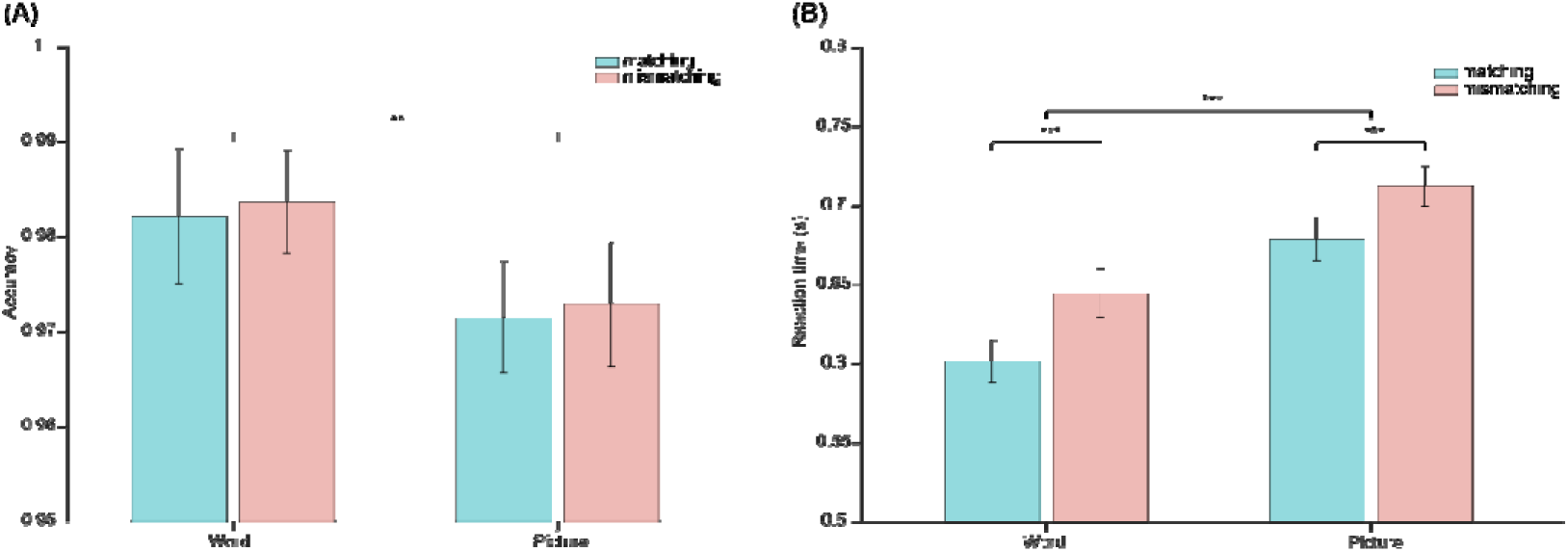
Bar plots of behavioral accuracy (a) and reaction time (b) for each condition.

Statistical analysis results for RT (see Figure 2b) showed a main effect for both visual format [F_(1,26)_ =229.43, *p* < 0.001, *η_p_*^2^ = 0.90] and matching [F_(1,26)_ = 22.82, *p* < 0.001, *η_p_*^2^ = 0.47] factors. Specifically, RTs were shorter in the matching condition compared to the mismatching condition, and shorter in the word visual format than the picture format. However, there was no significant interaction effect between the two factors [F_(1,26)_ =1.19, *p* = 0.29, *η_p_*^2^ = 0.044].

### 3.2 Time-resolved decoding

Figure 3a shows the time-resolved decoding results between matching and mismatching trials separately for words and pictures. Classification accuracy for distinguishing matching from mismatching trials rose significantly above chance from approximately 370 ms to 800 ms for words and from approximately 390 ms to 800 ms for pictures.

**Figure 3.**
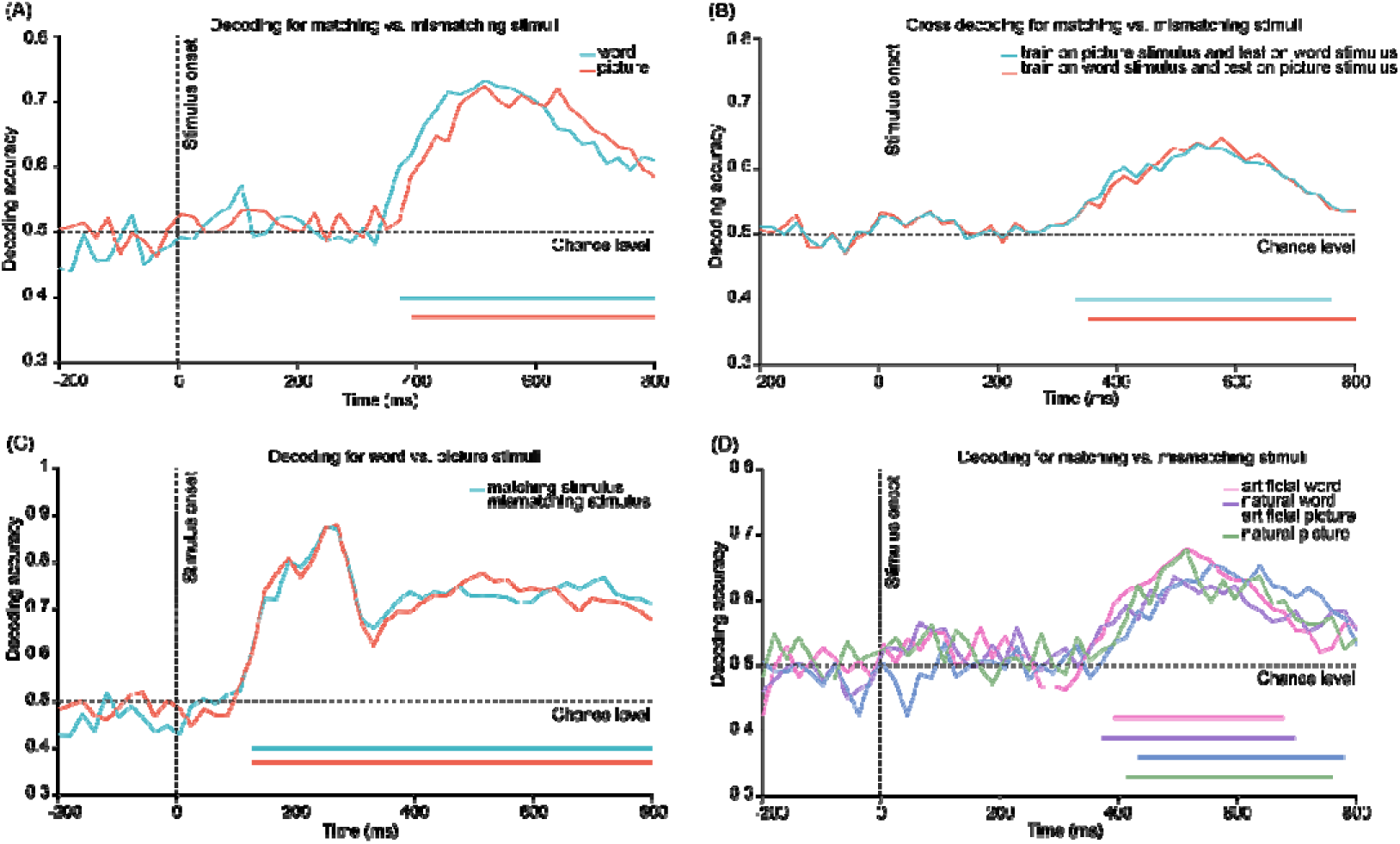
The decoding performance for four independent decoding analyses. (a) Within-condition decoding: match versus mismatch trials were decoded separately for the word and picture conditions. (b) Cross-condition decoding: classifiers were trained on one stimulus format (word or picture) and tested on the other to assess shared semantic representations across formats. (c) Visual format decoding: word versus picture trials were decoded separately for matching and mismatching conditions. (d) Category-specific decoding: match versus mismatch trials were decoded separately for natural and man-made objects within each stimulus format.

Similarly, the cross-decoding analysis (see Figure 3b) showed that discrimination between matching and mismatching trials emerged significantly earlier for written words (trained on pictures, from about 330ms) than for pictures (trained on words, from about 350ms).

Figure 3c shows the decoding accuracy between word and picture formats for matching and mismatching trials separately. Differences between word and picture categories emerged significantly above chance from approximately 125 ms to 800 ms in both conditions.

Figure 3d shows that, within the word visual format, significant differences between matching conditions emerged above chance level later for artificial objects (from approximately 390 ms) than for natural objects (from approximately 370 ms). Similarly, within the picture format, decoding accuracy for distinguishing matching from mismatching trials rose significantly above chance earlier for natural objects (from approximately 410 ms) than for artificial objects (from approximately 430 ms).

### 3.3 Time×time generalizations

Figure 4a presents the TGM for decoding between matching stimuli under the word condition. Decoding accuracy for word visual format rose significantly above chance from approximately 340 ms following stimulus onset. Significant decoding was nearly closed to the diagonal of the matrix, indicating highly time-specific neural representations and a predominantly dynamic coding pattern. Decoding accuracy for the picture visual format became significantly above chance from approximately 360 ms to 800 ms (Figure 4b). The TGM exhibited extensive off-diagonal generalization, indicating that neural representations generalized across multiple time points and were relatively stable over time.

**Figure 4.**
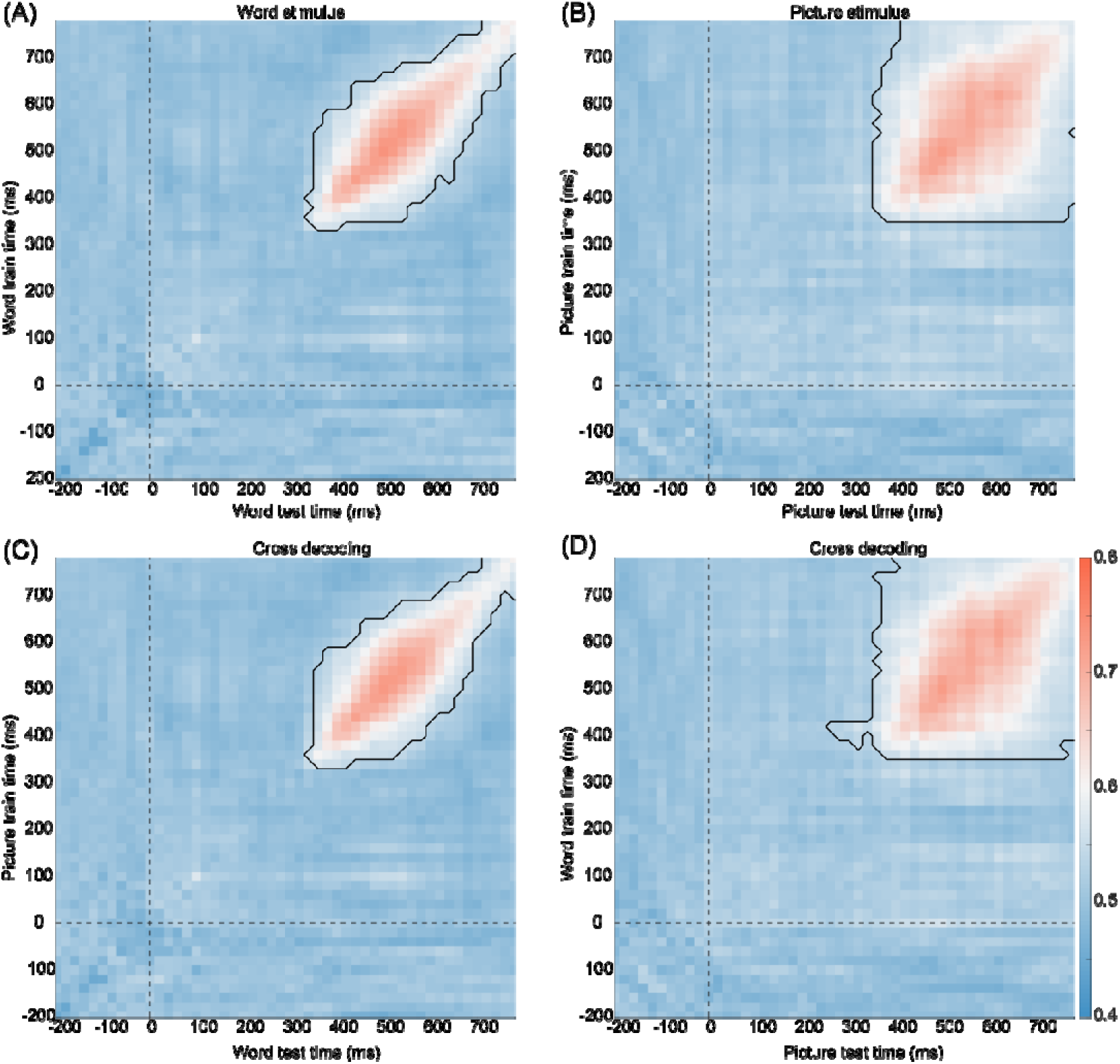
Temporal generalization analyses of semantic matching. (a-b) Within-condition temporal generalization matrices for decoding match versus mismatch trials in the word (a) and picture (b) conditions. (c-d) Cross-condition temporal generalization matrices obtained by training classifiers on one stimulus format (word or picture) and testing on the other. Black contours denote time points at which decoding accuracy was significantly above chance level (cluster-based permutation test, p < .05), indicating reliable above-chance decoding. Diagonal patterns indicate temporally specific representations, whereas off-diagonal generalization reflects representations that remain stable across time.

To determine whether these representations were shared across stimulus formats, we conducted cross-decoding analyses by training classifiers on one modality and testing them on the other. The resulting TGMs closely resembled the within-modality results. Specifically, classifiers trained on pictures and tested on words reproduced the temporal dynamics observed in the word condition (Figure 4c), whereas classifiers trained on words and tested on pictures reproduced the pattern observed in the picture condition (Figure 4d). These findings suggest that word and picture processing rely on partially shared neural representations despite differences in their temporal generalization profiles.

Figure 5 presents the cross-modal TGMs of word and picture formats separate for the matching and mismatching conditions. For the matching condition (Figure 5a), two significant clusters were identified: a off-diagonal cluster spanning approximately 120–320 ms and 120–800 ms, and a diagonal cluster extending from approximately 320 to 800 ms. For the mismatching condition (Figure 5b), a similar off-diagonal cluster was observed between approximately 120–300 ms and 120–620 ms. In addition, a broad rectangular cluster emerged from approximately 400 to 800 ms, indicating more sustained temporal generalization during later processing stages than in the matching condition.

**Figure 5.**
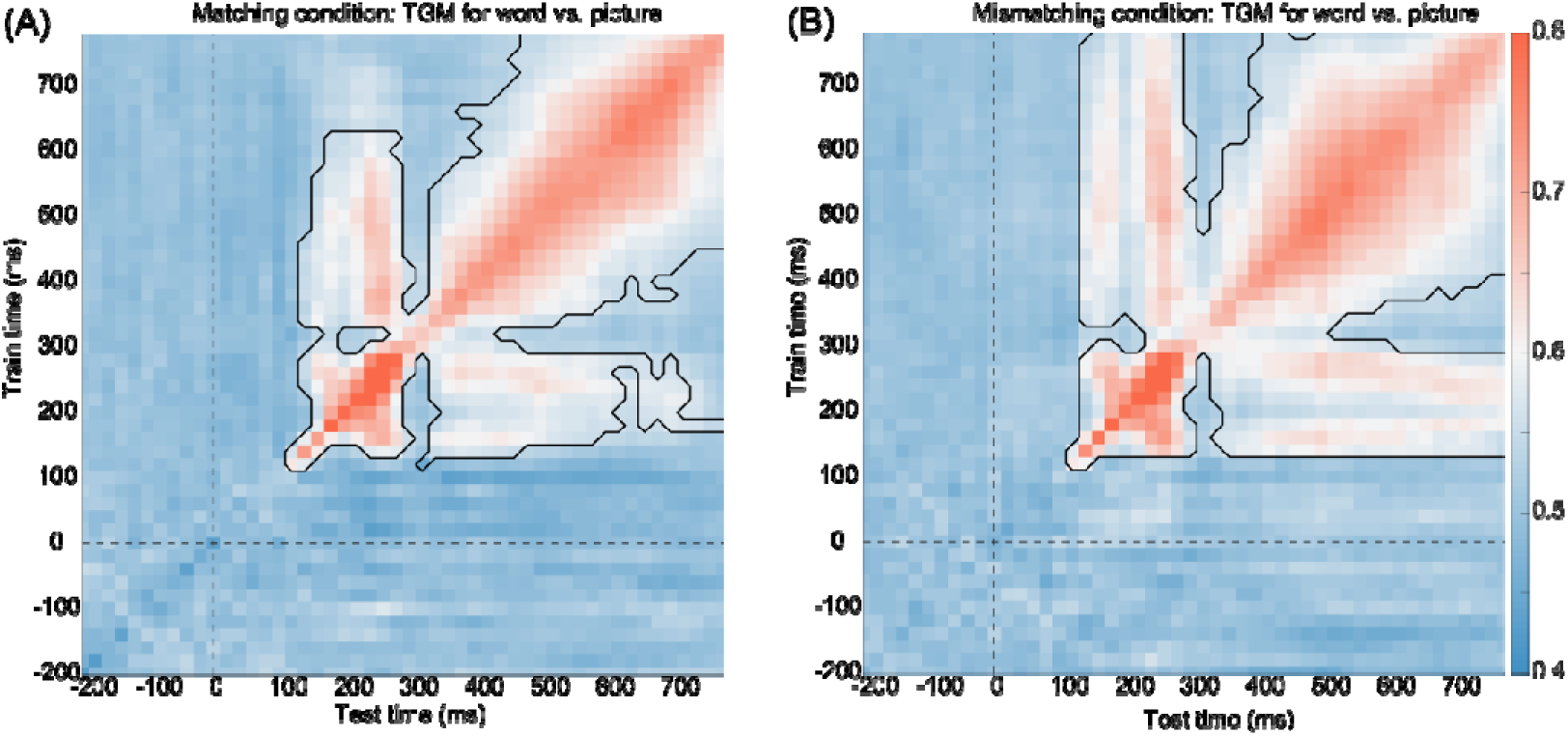
Visual format decoding results. Temporal generalization matrices for decoding word versus picture stimuli under match (a) and mismatch (b) conditions. Black contours indicate time points where decoding accuracy significantly exceeded chance level, based on cluster-based permutation testing (p < .05). Diagonal patterns indicate temporally specific representations, whereas off-diagonal generalization reflects representations that remain stable across time.

To assess format-related differences, temporal generalization analyses were performed separately for natural and artificial objects. Under the word condition (Figures 6a and 6b), both categories showed significant decoding primarily along the diagonal from approximately 360 to 680 ms, indicating largely time-specific representations. Under the picture condition (Figures 6c and 6d), significant decoding extended across a broad rectangular region between approximately 400 and 800 ms for both categories, reflecting substantial off-diagonal generalization. Thus, similar to the overall results, picture-based processing exhibited more temporally stable representations than word-based processing regardless of object category.

**Figure 6.**
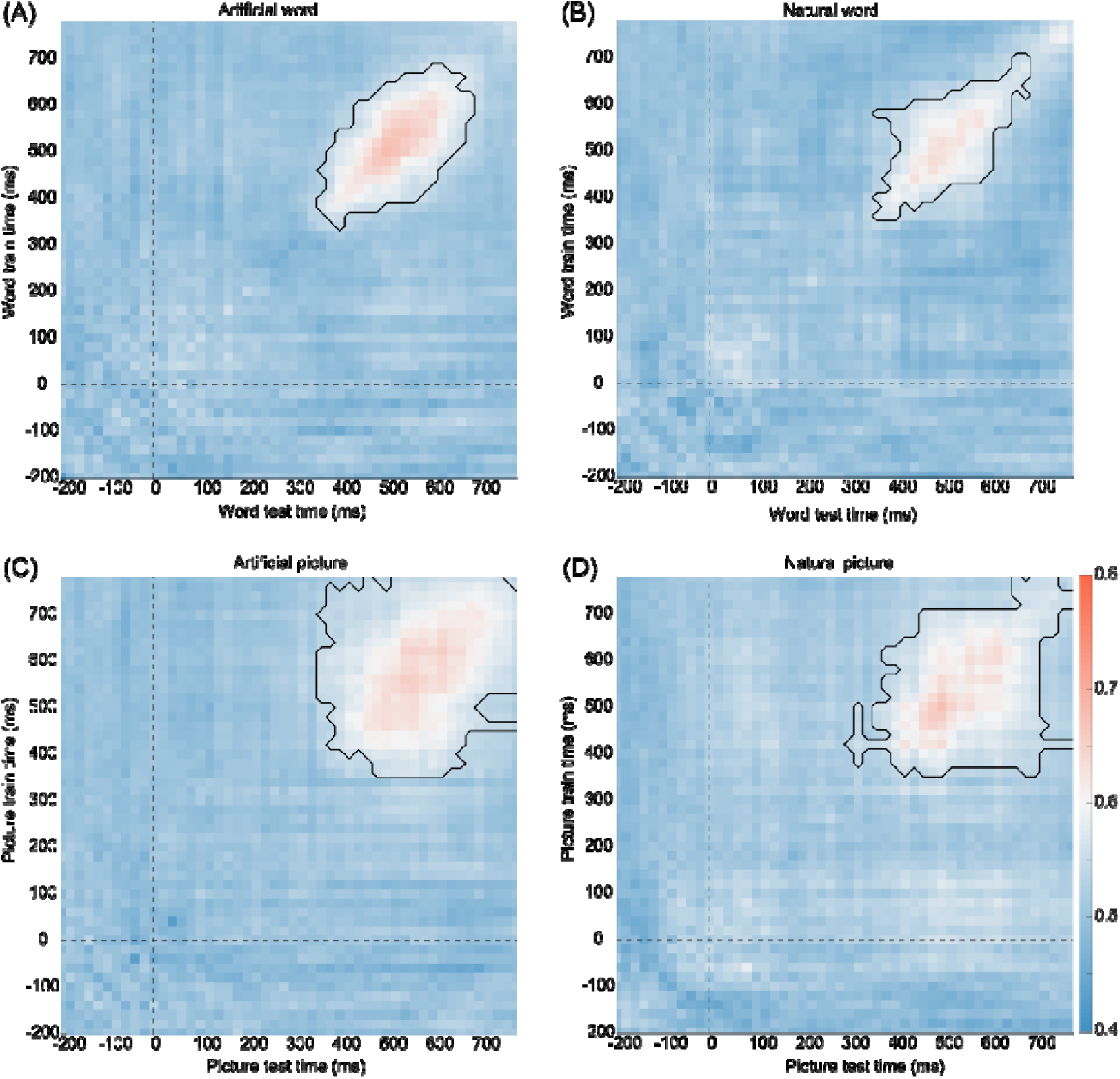
Category-specific temporal generalization decoding results. (a-d) Temporal generalization matrices for match versus mismatch decoding, separately for natural and man-made objects within the word and picture conditions. Black contours indicate significant above-chance decoding (cluster-based permutation test, p < .05). Diagonal patterns indicate temporally specific representations, whereas off-diagonal generalization reflects representations that remain stable across time.

## 4. Discussion

In the present study, we investigated the temporal dynamics of semantic matching between spoken words and visual information (i.e., written word or picture) using time-resolved decoding and temporal generalization methods. Several main findings emerged. First, reliable decoding of matching vs. mismatching trials was observed in both speech-word and speech-picture tasks. Second, matching-related decoding emerged earlier for written words than for pictures, suggesting earlier alignment between auditory linguistic input and visual word information than between auditory linguistic input and visual object information. Third, successful cross-decoding indicated partially shared congruency representations across formats, whereas direct decoding between words and pictures revealed robust format-specific information throughout processing. Fourth, temporal generalization analyses revealed distinct representational dynamics across formats. Finally, natural objects showed earlier semantic discrimination than artificial objects. These findings support the view that audiovisual matching relies on partially shared neural representations across representational formats, but that the timing and temporal evolution of auditory–visual alignment differ depending on visual format and object category.

### 4.1 Behavioral performance mirrors neural decoding dynamics

The behavioral findings generally converged with the decoding results (see the discussion from the following sections). Participants responded more accurately and faster in the word condition than in the picture condition, suggesting that semantic judgments were easier when information was presented as written words. This finding is broadly consistent with evidence that word and picture processing engage partially distinct routes to conceptual knowledge, with semantic information becoming available at different stages across modalities (Dirani & Pylkkänen, 2023). Whereas written words share linguistic representational codes with spoken words, pictures require mapping between auditory linguistic input and visual object representations before semantic congruency can be evaluated. Consequently, the longer response times observed for pictures likely reflect additional visual object processing and audiovisual integration demands, rather than slower semantic access per se (Ghuman et al., 2026).

Reaction times were also shorter for matching than mismatching trials, indicating a behavioral advantage for semantically congruent information. Such congruency facilitation effects are consistent with findings showing that semantically consistent information is integrated more efficiently, whereas incongruent information requires prolonged semantic evaluation and integration processes (Krugliak et al., 2024). The absence of a significant Format×Matching interaction indicates that the magnitude of the semantic congruency benefit was statistically comparable across word and picture conditions at the behavioral level. This pattern is consistent with evidence that words and pictures ultimately engage shared conceptual representations despite differences in their processing routes and temporal dynamics (Dirani & Pylkkänen, 2023, 2024). Thus, semantic congruency may exert its facilitative influence on a common conceptual system regardless of input format, even though the neural processes supporting auditory–visual alignment and integration differ across modalities.

### 4.2 Earlier semantic matching in the speech-word condition: the role of cross-modal representational alignment

A key finding of the present study is that matching-related information became decodable earlier in the speech-word condition than in the speech-picture condition. We interpret this temporal advantage as reflecting more efficient alignment between auditory word representations and visual word information, rather than faster semantic access for written words per se (Ghuman et al., 2026; Ralph et al., 2017). Written words share linguistic representational codes with spoken words, whereas pictures require visual object information to be extracted and organized before it can be integrated with auditory linguistic input (Grill-Spector & Weiner, 2014; Martin, 2016). Thus, the delayed onset in the picture condition may reflect additional visual-to-auditory integration demands during semantic matching.

This interpretation aligns with dual coding theory (Bi, 2021), which proposes that words and pictures are processed via partially distinct representational systems (verbal vs. imagery-based systems). Recent extensions and empirical work further support this distinction, showing differential contributions of verbal and non-verbal codes to memory and semantic integration (Monzel et al., 2022). According to this framework, pictures typically engage richer perceptual and imaginal representations, which may require additional processing before they can be aligned with auditory linguistic input during semantic matching. In contrast, written words share representational characteristics with spoken language and may therefore facilitate more rapid audiovisual alignment.

Complementary support comes from hub-and-spoke models of semantic cognition, in which modality-specific perceptual systems (“spokes”) interact with transmodal conceptual representations supported by anterior temporal regions (Patterson et al., 2007; Ralph et al., 2017; Rogers et al., 2004). Within this framework, visual object recognition requires the transformation of perceptual features into conceptual object representations before these representations can be aligned with auditory linguistic input. This additional transformation stage may contribute to the delayed emergence of matching-related neural information in the picture condition.

### 4.3 Cross-decoding and partially shared semantic representations

The cross-decoding results demonstrated successful generalization across word and picture formats (see Figure 3b), indicating the presence of shared neural representations underlying semantic matching (Bezsudnova et al., 2024; Giari et al., 2020). Specifically, classifiers trained on one modality were able to decode match versus mismatch in the other modality, albeit with slightly shifted temporal profiles. This finding provides evidence that partially shared neural information contributes to semantic matching across both linguistic and visual input streams. Importantly, the successful cross-decoding observed between speech-word and speech-picture conditions suggests that both tasks ultimately converged on partially shared semantic representations, even though the timing of representational alignment differed across formats.

Such results are consistent with distributed theories of conceptual knowledge, which propose that semantic matching representations are not tied to a single input modality but instead emerge from interactions among multiple modality-specific systems. Successful cross-decoding therefore suggests that word and picture processing recruit partially overlapping neural codes during semantic matching.

Recent multivariate neuroimaging studies using cross-modal decoding and representational similarity analysis have similarly demonstrated that semantic categories and object identity can be decoded across visual and linguistic formats, supporting the idea of partially shared conceptual coding across modalities (e.g., Dirani & Pylkkänen, 2023, 2024; Giari et al., 2020; Guan et al., 2025; Leonardelli et al., 2019; Li Calzi et al., 2025). Importantly, the present results extend these findings by showing that such shared representations are not only modality-independent but also temporally dynamic, emerging within a rapidly evolving neural time window. Furthermore, successful cross-decoding coexisted with robust within-time decoding between word and picture formats (see Figure 3c), indicating that shared semantic information and format-specific information were simultaneously represented in the neural activity patterns.

Importantly, temporal generalization analyses across modalities (see Figure 5) further demonstrated that these shared representations were not restricted to isolated time points. Significant cross-modal generalization extended across broad temporal windows in both matching and mismatching conditions, indicating that neural information related to semantic matching could be sustained and transferred across representational formats (King & Dehaene, 2014). Notably, the mismatching condition showed a larger late-stage generalization cluster than the matching condition, suggesting that semantic conflict or incongruency may engage more prolonged representational processing. These findings extend the time-resolved decoding results by showing that shared semantic information is maintained over extended processing intervals rather than emerging only transiently.

### 4.4 Temporal generalization patterns: dynamic vs. stable representations

Temporal generalization analysis further revealed striking differences between word and picture processing. Word decoding was primarily closed to the diagonal of the TGM matrices, indicating that neural representations were highly time-specific and rapidly changing over time. In contrast, picture decoding exhibited broad off-diagonal generalization, suggesting more temporally stable and sustained neural representations (King & Dehaene, 2014). Such rapidly evolving representations are consistent with a sequence of processing stages that may include orthographic, lexical, and semantic operations.

This dissociation suggests that word processing may rely on rapid transitions among orthographic and linguistic representations that can be efficiently aligned with auditory input, resulting in neural patterns that evolve quickly over time (Hauk et al., 2006). In contrast, picture processing likely involves the gradual accumulation and integration of visual object information before stable auditory–visual matching representations can be established, giving rise to more sustained neural representations (Cichy et al., 2014; Contini et al., 2017; DiCarlo et al., 2012; Ghazaryan et al., 2023). Consequently, picture-based semantic matching may recruit more sustained neural representations, giving rise to the extensive off-diagonal generalization observed in the present study.

### 4.5 Natural vs. artificial objects: category-dependent temporal dynamics

We further found that natural objects elicited earlier decoding onset than artificial objects across both word and picture formats. These findings suggest that object category may modulate the speed with which visual information can be integrated with auditory linguistic input during semantic matching. One possible explanation is that natural objects (e.g., fruits and animals) possess more perceptually coherent and highly shared feature structures, whereas artificial objects are more strongly defined by functional and associative properties (Hebart et al., 2020; Ralph et al., 2017). Such differences may contribute to the later emergence of semantic discrimination observed for artificial objects.

This interpretation is consistent with contemporary semantic cognition frameworks, which propose that conceptual representations emerge from interactions between modality-specific perceptual systems and transmodal semantic hubs, with natural categories relying relatively more on perceptual feature statistics and artifact categories relying more heavily on functional knowledge and contextual associations (Ralph et al., 2017). Furthermore, recent EEG decoding and representational similarity studies have shown that object-category information emerges rapidly during visual processing and that neural representational geometry is systematically shaped by semantic category structure (Hebart et al., 2023; Iamshchinina et al., 2022; Wamain et al., 2023; Weber et al., 2024). These representational properties may contribute to the earlier decoding onset observed for natural objects in the present study.

### 4.6 Limitations

Several limitations should be considered when interpreting the present findings. First, the study focused on healthy young adults using a controlled EEG paradigm, which may limit generalizability to other populations such as older adults, clinical groups, or developmental samples (Marsicano et al., 2024; Ng et al., 2022; Turoman et al., 2024). Second, although time-resolved MVPA and temporal generalization analyses provide strong temporal resolution, EEG inherently lacks precise spatial resolution, making it difficult to localize the exact neural sources underlying the observed effects. Future studies combining EEG with fMRI or MEG could help resolve this limitation (Peelen & Downing, 2023).

An additional limitation concerns the interpretation of the earlier decoding onset observed for written words. Because the present paradigm required audiovisual semantic matching, the observed latency differences may reflect not only semantic access but also modality-specific integration processes between auditory and visual information. Future studies directly comparing unimodal word and picture processing with audiovisual matching paradigms will be necessary to determine the extent to which the present effects reflect representational alignment and audiovisual integration processes, as opposed to differences in semantic access per se.

Fourth, the present analyses primarily relied on linear classification approaches (e.g., linear SVM) and time-resolved decoding. While this approach is widely used in cognitive neuroscience, it may not capture potentially nonlinear dynamics of semantic processing (Hosseini et al., 2021). Future work could explore deep learning-based decoding models or nonlinear representational analyses to further characterize semantic dynamics (Craik et al., 2019).

Finally, although the present paradigm was designed to approximate cross-modal semantic matching, it remains considerably more constrained than everyday language comprehension. Traditional trial-based and forced-choice paradigms provide strong experimental control but may not fully capture the continuous and dynamic nature of real-world language processing (Alday, 2019; Brodbeck & Simon, 2020). Moreover, explicit matching judgments likely recruit additional attentional, response-selection, and decision-making mechanisms that can contribute to late positive neural activity beyond semantic processing itself (S. Luck, 2014; Polich, 2007). Future studies should therefore examine semantic matching under more naturalistic conditions, such as continuous speech comprehension, naturalistic narratives, or passive viewing paradigms, which may better reflect how conceptual information is processed in everyday communication (Alday, 2019; Welke & Vessel, 2022).

## CRediT authorship contribution statement

**Ying Xin**:Data Curation, Writing – review & editing, Writing – original draft, Visualization, Validation, Software, Resources, Methodology, Investigation; **Huashuai Xu**: Writing – review & editing, Writing – original draft, Methodology; **Fengyu Cong**:Writing – review & editing, Writing – original draft,Visualization; **Weiqi He**: Writing – review & editing, Writing – original draft, Visualization, Validation, Supervision; **Guanghui Zhang**:Writing – review & editing, Writing – original draft, Visualization, Validation, Supervision, Software, Resources, Methodology, Conceptualization;

## Data availability

The data and analysis scripts are available from the corresponding author upon reasonable request.

## Acknowledgments

We gratefully acknowledge the assistance of Xinran Wang in data collection, and thank all participants for their time, effort, and cooperation during the experiment.

